# A cell-based probabilistic approach unveils the concerted action of miRNAs

**DOI:** 10.1101/298596

**Authors:** Shelly Mahlab-Aviv, Nathan Linial, Michal Linial

**Affiliations:** The Rachel and Selim Benin School of Computer Science and Engineering, The Hebrew University of Jerusalem, Jerusalem, Israel; Department of Biological Chemistry, Institute of Life Sciences, The Hebrew University of Jerusalem, Jerusalem, Israel

**Keywords:** Actinomycin D, CLIP-Seq, miRNA-target prediction, Stochastic model, TargetScan, ceRNA, miRNA binding sites, miRbase, Cell simulation, Markov chain, human cell-line

## Abstract

Mature microRNAs (miRNAs) regulate most human genes through direct base-pairing with mRNAs. We investigate some underlying principles of such regulation. To this end, we overexpressed miRNAs in different cell types and measured the mRNA decay rate under transcriptional arrest. Parameters extracted from these experiments were incorporated into a computational stochastic framework which was developed to simulate the cooperative action of miRNAs in living cells. We identified gene sets that exhibit coordinated behavior with respect to all major miRNAs, over a broad range of overexpression levels. While a small set of genes is highly sensitive to miRNA manipulations, about 180 genes are insensitive to miRNA manipulations as measured by their degree of mRNA retention. The insensitive genes are associated with the translation machinery. We conclude that the stochastic nature of miRNAs reveals an unexpected robustness of gene expression in living cells. Moreover, the use of a systematic probabilistic approach exposes design principles of cells’ states and in particular, the translational machinery.

**Highlights:**

- A probabilistic-based simulator assesses the cellular response to thousands of miRNA overexpression manipulations
- The translational machinery displays an exceptional resistance to manipulations of miRNAs.
- The insensitivity of the translation machinery to miRNA manipulations is shared by different cell types
- The composition of the most abundant miRNAs dominates cell identity

## Introduction

Mature microRNAs (miRNAs) are small, non-coding RNA molecules (~22 nucleotides) that regulate genes through base-pairing with their cognate mRNAs, mostly at the 3′ untranslated region (3’-UTR) (Ameres and Zamore, 2013; Moore et al., 2015; Pasquinelli, 2012). In multicellular organisms, miRNAs act post-transcriptionally by affecting the destabilization and degradation of mRNAs, as well as interfering with the translation machinery (Chekulaeva and Filipowicz, 2009; Eichhorn et al., 2014; Filipowicz et al., 2008). Switching between cell states is accompanied by a shift in the profile of miRNAs (Pelaez and Carthew, 2012). Indeed, miRNA-dependent transitions are documented in cells undergo quiescence (Cheung et al., 2012), differentiation (Yang et al., 2013), viral infection (Zhang et al., 2012) and cancer transformation (Bertoli et al., 2015; Lu et al., 2005).

In humans, there are ~2500 mature miRNAs that derive from ~1900 genes (Ameres and Zamore, 2013). Studies of miRNA-mRNA regulatory networks reveal that almost all coding genes have multiple putative miRNA binding sites (MBS) at their 3’-UTR (Landgraf et al., 2007; Liang et al., 2007; Stark et al., 2005), and many miRNAs can possibly target hundreds of transcripts (Balaga et al., 2012; Rajewsky, 2006). However, current estimates postulate that only ~60% of the human coding genes are regulated by miRNAs (Ha and Kim, 2014; Jonas and Izaurralde, 2015). Most of our knowledge of the specificity of miRNA-mRNA network is based on computational prediction tools (Peterson et al., 2014) that use parameters learned from in-vitro overexpression or miRNA knockdown experiments (Hausser and Zavolan, 2014). The output of the CLIP-Seq methodologies is a collection of interactions between miRNAs and mRNAs from healthy and diseased cells (Li et al., 2014). The results from a revised CLIP-seq protocol, called CLASH, are direct interactions of miRNAs with their binding sites sequences (MBS) on mRNAs (Helwak et al., 2013; Moore et al., 2015). Unfortunately, many of the above protocols suffer from low coverage and poor consistency (discussed in (Chi et al., 2012)).

A quantitative perspective for miRNAs regulation is strongly dependent on the identity and quantity of limiting factors in the cell such as the AGO protein, which is the catalytic component of the RNA silencing complex (RISC) (Janas et al., 2012; Wen et al., 2011). From the mRNA perspective, the number of miRNA molecules, and the positions of MBS along the relevant transcript determine the potential of miRNA interactions (Agarwal et al., 2015). The outcome is a rich regulatory network displaying a “many to many” relation of miRNAs and mRNAs. Such design supports noise reduction (Ebert and Sharp, 2012; Herranz and Cohen, 2010; Schmiedel et al., 2015), and robustness against environmental fluctuation (Li et al., 2009).

A cellular view of miRNAs network was formulated by the ceRNA hypothesis (Denzler et al., 2014; Salmena et al., 2011). Accordingly, an overexpression of MBS-rich molecules of RNA may displace miRNAs from their primary authentic targets (Denzler et al., 2016; Tay et al., 2014), resulting in an attenuation relief of specific mRNAs. The result of such a competition is an interplay between direct and indirect effects on gene expression (Seok et al., 2016; Yuan et al., 2015). The dynamics of the miRNA-target regulatory network in view of direct and distal regulation had been modeled (Nitzan et al., 2014). It was further postulated that many of the miRNA weak sites contribute to target-site competition without imparting repression (Denzler et al., 2016).

In this paper, we describe a quantitative stochastic model that challenges the cell steady-state in view of alteration in miRNAs’ abundance. The model operates at the cellular level and compares the overall trend of miRNA regulation in various cell lines. We systematically analyzed the behavior of miRNA-mRNA regulation in various cell types. We confirm that the stochastic nature of miRNA regulation reveals an unexpected robustness of the miRNA regulation in living cells.

## Results

### Determining miRNAs stability and decay rate of mRNAs upon transcriptional arrest

The nature and extent of miRNA regulation in living cells are depicted by the absolute quantities, composition, and stoichiometry of the miRNAs and mRNAs (Arvey et al., 2010). In this study, we model the outcome of the miRNA-mRNA network under a simplified paradigm where synthesis of new transcripts (miRNA and mRNAs) is prevented.

We first tested the relative changes in the quantities of miRNAs and mRNAs in HeLa and HEK-293 cell-lines, in the presence of the transcriptional inhibitor Actinomycin D (ActD, Figure 1A). Overall, we mapped 539 and 594 different miRNAs in untreated HeLa and HEK-293 cells, respectively (Figure 1A). In addition, 16,236 and 16,463 different expressed mRNAs (not including miRNAs) were mapped from HeLa (Supplemental Dataset S1) and HEK-293 cells (Supplemental Dataset S2), prior to ActD treatment, respectively. We then tested the composition of miRNAs and mRNAs 24 hrs post-treatment. Importantly, the number of miRNA molecules 24 hrs after the application of the drug remains constant in HeLa (Spearman rank correlation, r = 0.94) and HEK-293 cells (Spearman rank correlation, r = 0.97, Figure 1B). In contrast, the number of mRNAs molecules has monotonically declined in accordance with the effect of ActD on the bulk of short-lived mRNAs (Figure 1A). Maximal variability in the profile of mRNAs is measured between 0 hr and 24 hrs for HeLa (Figure 1B, Spearman rank correlation, r = 0.84, top right) and HEK-293 cells (Spearman rank correlation, r = 0.88, bottom right, right). Supplemental Figures S1 and S2 show the pairs for all other time points for HeLa and HEK-293, respectively.

**Figure 1.**
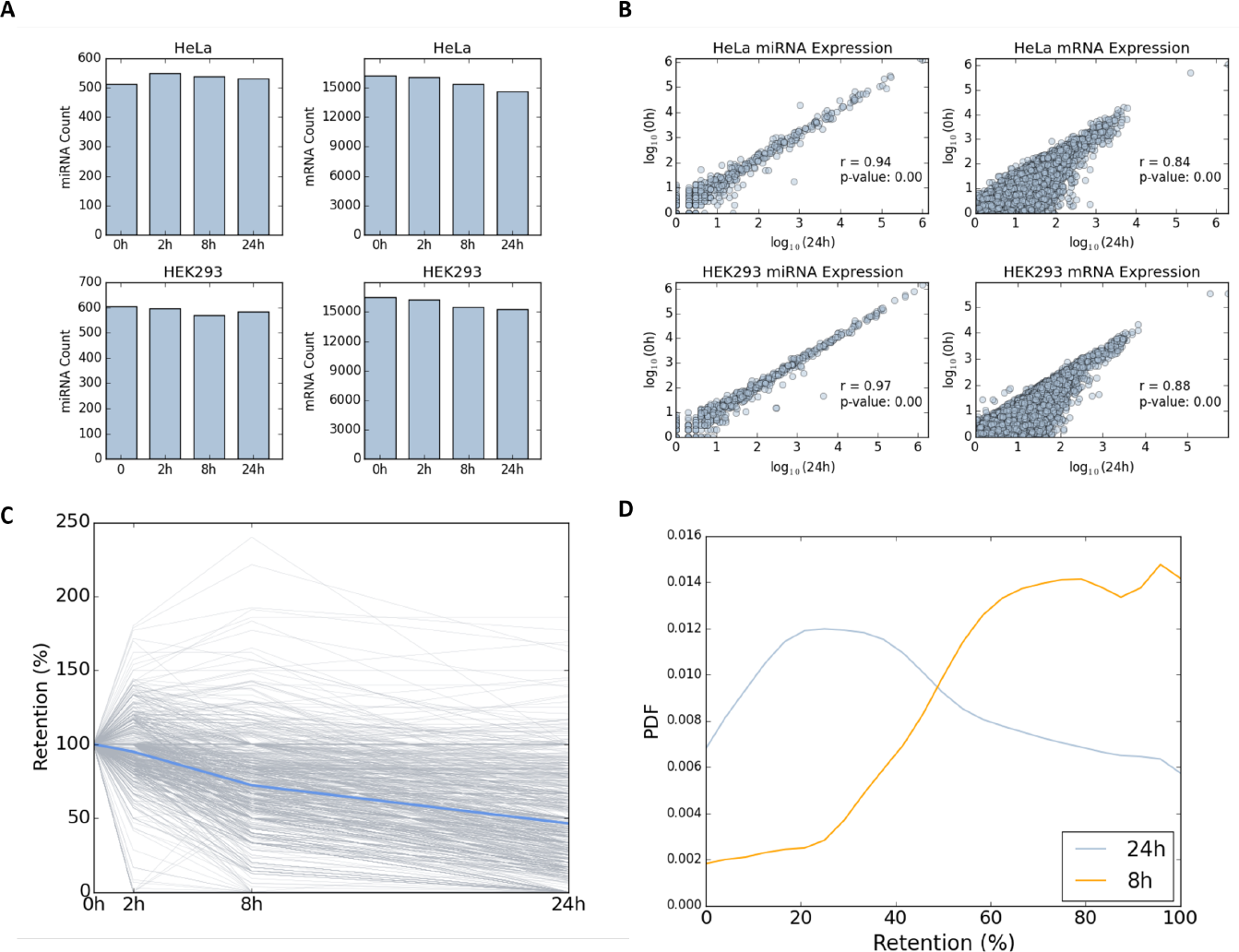
Expression profiles of miRNA and mRNA under transcription arrest. Counting of miRNA miRNAs (left) and mRNAs (right) for the 4 different time points for HeLa (top) and HEK-293 (bottom). **(A)** The samples were collected at 0, 2, 8, and 24 hrs following transcription inhibition by ActD. **(B)** Expression of miRNAs (left) and mRNAs (right) in pairs of 4 different time points for HeLa (top) and HEK-293 (bottom). Gene expression is presented by logarithmic scale (log_10_). Spearman correlation (r) is listed for each pair along with the p-value of the significance. Source data is available in Supplemental Dataset S1 (HeLa) and Supplemental Dataset S2 (HEK-293). **(C)** Relative abundance of each expressed mRNAs for HeLa cells. At time 0, the relative abundance is set to 100%, and at each proceeding time points the abundance relative to time 0 is reported. Each line represents a single gene (mRNA). Only genes with a minimal expression level of 0.02% expression are listed (equivalent to 97 FPKM, total of 860 genes). The blue line represents the average of all reported genes at each time point. **(D)** Compilation of mRNA retention distribution (PDF) of all the reported genes after 8 hrs and 24 hrs from initiation of transcription inhibition by ActD. All genes with a retention level ≥100 are combined (at 100% retention).

Figure 1C follows the change in the expression level of individual genes in HeLa cells along 24 hrs from ActD treatment. Changes in each mRNA abundance were quantified relative to the abundance at the starting point. To secure the stability of the results, we only report on the retention percentage for genes that are expressed above a predetermined threshold (total 860 genes, Figure 1C). We illustrate how the distribution of the retention level (in %, 860 genes) varies between two different time points (Figure 1D). The average retention rate 8 hrs after ActD treatment is ~83% and decreases to 53% after 24 hrs. These results validate that the decay rate for most mRNAs is a gradual process that continues for 24 hrs.

### Direct targets and non-target mRNAs are affected by miRNAs overexpression

Figure 2 shows the results of direct and indirect effects of overexpressing hsa-mir-155, a representative of expressing miRNA. HeLa cells were transfected with individual miRNAs, and the number of miRNA and mRNA molecules were measured 24 hrs after the introduction of ActD. We quantified the effect of hsa-mir-155 by considering its targets. Specifically, for each miRNA, we split all expressed genes to two sets - targets and non-targets, according to the TargetScan 7.1 prediction table (Agarwal et al., 2015) (see Materials and Methods). Retention rates of all genes relative to their starting point are shown (Figure 2A). The average decay rate for hsa-mir-155 direct targets is slightly faster compared with the set of the non-targets (Figure 2A, compare pink and blue thick lines). Furthermore, the decay after 24 hrs of ActD treatment for HeLa cells overexpressing hsa-mir-155 is enhanced in the transfected vs. naïve cells (Figure 2B, upper panel). While the shift in the relative mean statistics (Figure 2B) for the direct targets is marginal (p-value = 0.12), the shift for the non-target genes is highly significant (p-value of 0.002, Figure 2B, compare solid and dashed lines). The result implies a certain degree of stabilization for the non-target genes as a result of hsa-mir-155 overexpression. A similar retention profile was observed by overexpressing hsa-mir-124a in HeLa cells (Supplemental Figure S3).

**Figure 2.**
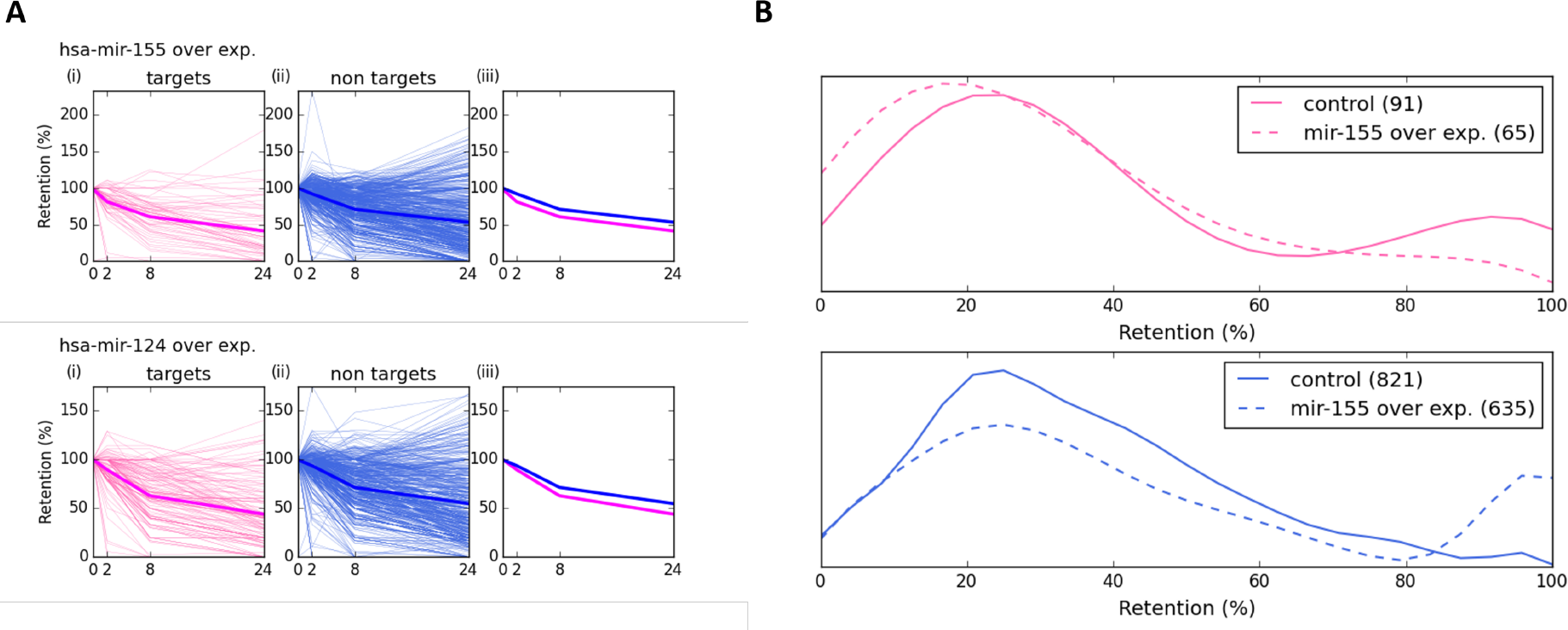
Retention profile of mRNAs following overexpressing miRNAs in HeLa cells. Relative mRNA retention in HeLa cells that were transfected and overexpressed with hsa-mir-155. Measurement were taken at 4 time points as indicated. **(A).** The retention plots are partitioned to target genes (i) (pink, left panels) and (ii) non-target genes (blue, middle panels). In (iii), the pink and blue lines are the average retention patterns for of hsa-mir-155 for targets and non-targets, respectively. **(B)** Distribution of genes retention after 24 hrs from ActD treatment, according to their labels as targets (upper panel, pink) and non-targets (lower panel, blue). The plots compare the retention of genes from the control (smooth line), and from hsa-mir-155 overexpressed condition (dashed line). The number of genes that are included in the analyses are indicated in parentheses. Target genes are marked by pink lines (top) and the non-target genes by blue lines (bottom). Note the shift in the distribution in the non-target genes towards the genes with higher retention level. All genes with a retention level ≥100 are shown as 100% retention.

We conclude that under the described experimental settings, the miRNA regulatory network affects the probabilities of miRNA-target interactions, mostly by an indirect propagation, presumably due to a competition on MBS, along with a continuous change in the miRNAs-mRNAs stoichiometry.

### Assessing a probabilistic approach for miRNA - mRNA interactions

The experimental results (Figures 1-2) emphasize the need for a systematic approach for studying miRNA-mRNA interaction network acting under the quantitative constraints of living cells. For our computational approach, we consider a stochastic process in which a dynamic competition between miRNAs and the available mRNAs occurs. In such a setting, the miRNA-mRNA binding probabilities dictate the effectiveness of gene expression attenuation. We used the TargetScan matrix of miRNA-MBS interactions, in which each miRNA-MBS interaction is associated with a probabilistic score. Such a score can be considered the probability of effective binding for any specific pair. Altogether TargetScan matrix reports on over 7M interactions. Importantly, genes that lack MBS in their 3’-UTR are not listed in TargetScan matrix. Altogether, there are 1,183,166 high quality pairs between miRNA and MBS that are considered in our study (see Materials and Methods).

We set to investigate the properties of the miRNA-mRNA interaction network in living cells. To this end, we developed an iterative simulator called COMICS (COmpetition of MiRNAs Interactions in Cellular Systems). Figure 3A illustrates a single iterative cycle of COMICS. The probabilistic framework relies on a constant update of the cell-state which is defined by the amounts and the distributions of miRNAs and mRNA types and the partition of occupied MBS and free molecules. COMICS iterations capture the stochastic process that characterizes miRNA regulation.

**Figure 3.**
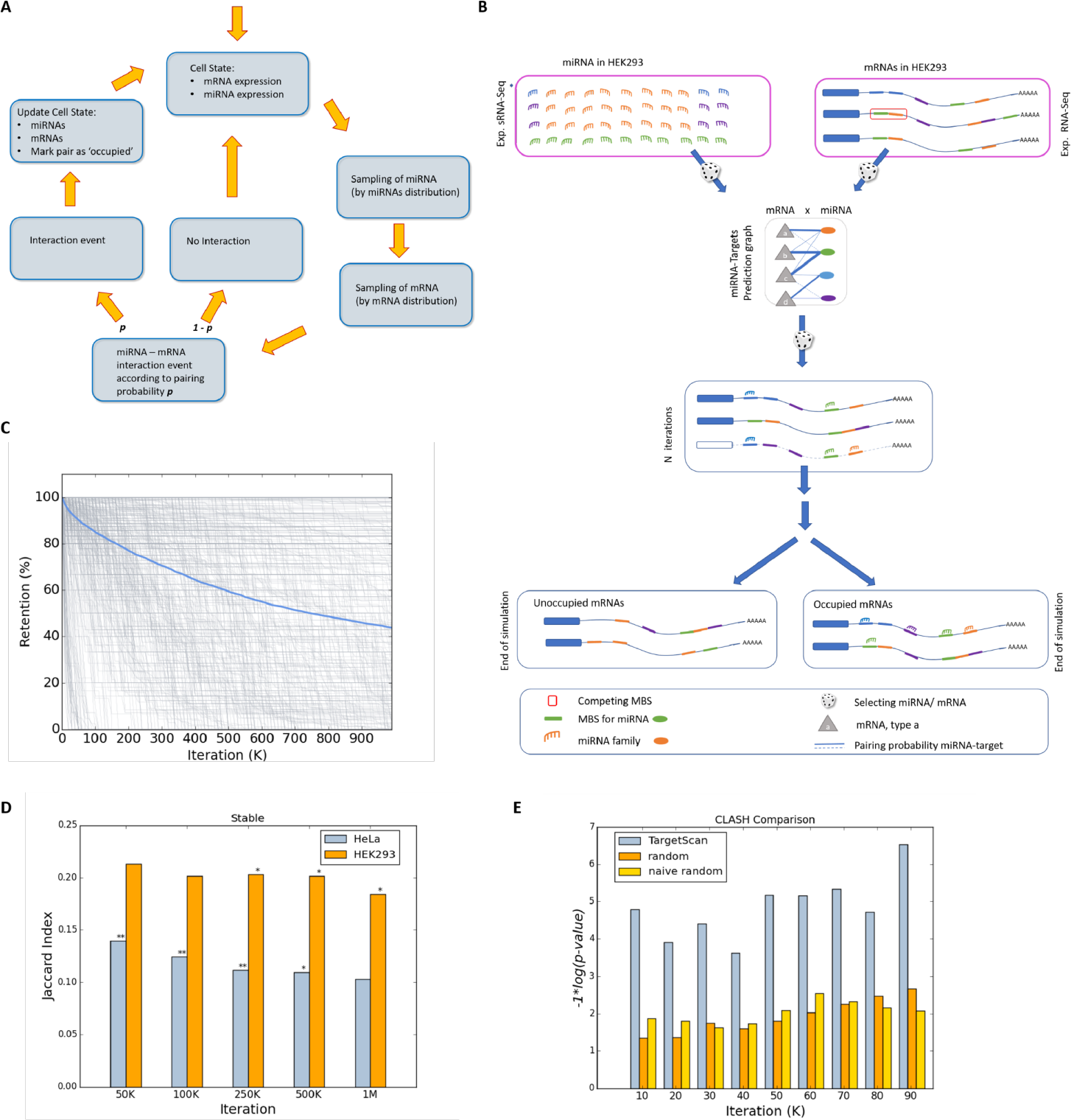
Scheme of the COMICS platform and performance of the simulation process. A schematic view for a single iteration of COMICS simulation. **(A)** A scheme for a single iteration step. After each successful interaction step, the distributions of the miRNA and mRNA in the cells are updated. Therefore, the next iteration is slightly changed due to resetting of the mRNA composition and availability of the free pool of miRNAs. The input for the simulator matches the molecular profile of the cell type under study. **(B)** The outline of the major steps of COMICS operation from the mRNA perspective. The composition of miRNAs in the cells is obtained from the experimental measurement at 0 hr), normalized for 50k miRNAs and of 25k mRNAs. For HeLa cells, these are 3666 types of mRNAs that are included in the analysis (i.e. above a minimal threshold). Sampling of the miRNA and mRNAs is done according to their distribution and the probability of the interaction according to TargetScan MBS interaction scores (with 1.2 M values). A dashed mRNA shown after N iterations signifies an occupied transcript that is still halted prior to its degradation, and releasing the bound miRNAs. **(C)** The retention of HeLa expressed genes along running COMICS simulations for 1M iterations. COMICS simulation on input from HeLa cells reports on 3666 types of mRNAs and 110 of miRNAs. These numbers account for the 50k and 25k molecules of miRNAs and mRNAs, respectively. Each grey line represents the retention profile of a single type on mRNA. The blue line shows the mean retention. For graphical clarity, only mRNA above a predefined expression (>0.02%) are shown. **(D)** Validation of COMICS performance in view of the results from transcription arrest in HeLa (grey) and HEK-293 cells (orange). At each of the indicated steps of the COMICS simulation run, the overlap in gene retention for the set of genes that remain stable (defined as >85% retention) was measured by calculating the Jaccard score. The statistical significance associated with the correspondence of the results (measured by p-value of the Fisher exact test) are indicated by asterisks * <0.05 and **, <0.005. **(E)** Testing COMICS performance and dependency on the information in TargetScan interaction matrix. COMICS simulation performance in HEK-293 was compared to the bounded pairs as reported from CLASH data on HEK-293. The histogram shows the performance in term of the significant of the overlap of the reported COMICS results (100k iterations) using TargetScan probabilistic converted matrix (grey), and two versions of randomization for the interaction table (Supplementary File S1, Supplementary Figure S5). The statistical test was based on the 251 genes that are reported as pairs miRNA-mRNA pairs by CLASH and expressed above the minimal expression threshold used for COMICS simulation protocol. The use of the TargetScan matrix shows significant results versus CLASH data (at the significant range p-value of 1e-4 to 1e-6). Applying any of the randomizations for the miRNA-MBS interaction table, caused a drop in the performance to non-significant values.

Figure 3B is a breakdown of the COMICS process through the lens of the regulated mRNAs. Specifically, the sampling process is driven by the composition of miRNAs and mRNAs and their absolute numbers in a cell (Figure 3B, pink frames). Recall that the expression profiles of miRNAs and mRNAs are a cell-specific (Supplemental Dataset S1 (HeLa); Dataset S2 (HEK-293)). Each mRNA is characterized by the types and positioning of its MBS at the 3’-UTR of the transcript. The interaction prediction table is associated with a probability-based score for any specific pairs of miRNA and MBS. In each iteration, a miRNA is sampled randomly, according to its relative abundance. Next, one of its target genes is chosen randomly according to the measured expressed mRNAs distribution. A binding event may occur according to each pair binding probability. Following such an event, the distribution of both, the miRNAs and mRNAs is updated (Figures 3A-3B). The status of the mRNA following a successful pairing is changed (i.e., marked as ‘prone to degradation’). The status of mRNA as ‘occupied’ does not prevent it from engaging in a subsequent binding, unless its other MBS are located in close proximity. Such overlapping MBS are defined according to a minimal spacing between them. The occupied mRNA is marked for degradation with some delay, reproducing the likely scenario of a cooperative binding by multiple miRNAs prior to degradation in vivo. Based on the validated stability of miRNAs (Figure 1B), once the occupied mRNA is removed, all bounded miRNAs return to the free miRNA pool. As a result, the stoichiometry of miRNA to mRNA is gradually changing with an increase in the ratio of miRNAs to free mRNAs.

Figure 3C shows the result of COMICS following one million iterations on HeLa cells. Note that following 1M iterations the mean retention of mRNAs is 43.5%, similar to the decay rate observed in living cells (Figures 1-2). Figure 3C shows the decay rate of 755 genes that are above a predetermined expression threshold (>0.02% of mRNA molecules). Supplemental Dataset S3 reports the stepwise output of mRNA levels along the 1M iteration run.

COMICS output and the actual cell experiments were compared by two complementary tests: (i) Calculating the correlation between unoccupied genes (i.e., genes that remain available following 24 hrs of ActD treatment, Figure 3D). (ii) Scoring the correspondence of genes that were occupied along COMICS iterative run (Figure 3E) and their correspondence with the results from CLASH experiment (Helwak et al., 2013) that was performed on HEK-293 cells. Figure 3D shows the results of comparing COMICS performance with experimental results considering high retention values (>85%). There are 122 such genes in HeLa and 158 genes in HEK-293. The hypergeometric test shows high statistical significance (100k iterations, HeLa cells, p-value = 0.00064). Similarly, the overlap with the results presented by CLASH (Helwak et al., 2013) remains highly significant throughout the iteration run (Figure 3E, hypergeometric test p-value = 0.0014). Replacing the TargetScan miRNA-MBS interaction table with two randomization modes (see Materials and Methods, and Supplemental File S1) completely eliminated the significant correspondence of the CLASH results and those of COMICS (Figure 3E). We conclude that the stochastic probabilistic protocol used by COMICS faithfully simulates miRNA regulation that takes place in living cells.

### Simulating miRNA overexpression by COMICS reveals stabilization of non-target genes

COMICS system is used to perform exhaustive overexpression experiments, while explicitly testing the effect of miRNA-mRNA probabilities on the mRNA decay. We activated COMICS by manipulating the abundance of hsa-mir-155 from its native state (x1, no overexpression) to varying degree of overexpression (x0.5, x3, x9, x18, x90, x300 and x1000) while respecting the probabilistic framework of the in-silico simulation. In practical terms, the overexpression of a single miRNA causes a change in the distribution of all other expressed miRNAs (Figure 4A). Figure 4A shows that following an overexpression by factors of x300 and x1000, as many as 20% and 50% of all miRNAs consist of hsa-mir-155 molecules, respectively. It is important to note that the calculated fraction of miRNAs in the cell following overexpression is a direct reflection of its abundance in the native cell. These miRNA profiles are strong characteristics of different cell types.

**Figure 4.**
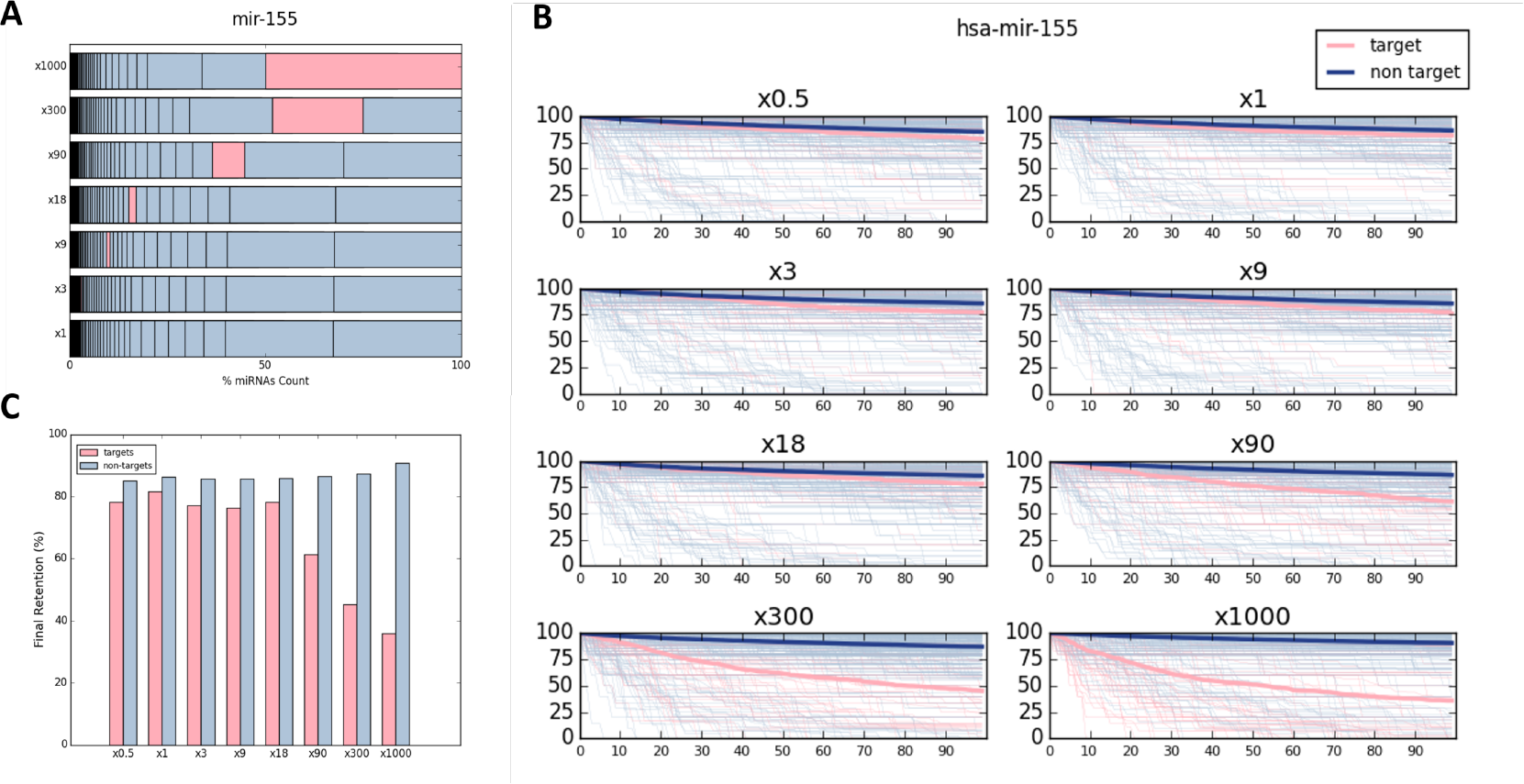
miRNA overexpression paradigm using COMICS platform. **(A)** The percentage of miRNA is sorted from lowest (left) to highest (right). Seven different hsa-mir-155 over expression simulations are shown (x1, x3, x9, x18, x90, x300 and x1000) from bottom to top. The percentage of hsa-mir-155 is marked in pink. The relative percentage of each miRNA abundance is separated by vertical line. **(B)** Overexpression simulation of hsa-mir-155 in eight different overexpressing factors, where hsa-mir-155 target genes are marked in pink and all other non-target genes are marked in gray. The mean retentions of both gene groups are plotted in bold lines. **(C)** Average final retention of the different simulation runs, using different overexpression factors of hsa-mir-155 as shown in B, of hsa-mir-155 target and non-target genes.

Figure 4B demonstrates the gradual change in mRNA retention of each gene (above a predetermined threshold) along 100k iterations of COMICS simulation (source data in Supplemental Dataset S4 for HeLa and HEK-293 cells). Figure 5B shows the gradual alteration in the overall dynamic of the gene retention following the increase in overexpression levels (according to the multiplication factors). We show that the final retention level is sensitive to the degree of overexpression (Figure 4C). In the case of hsa-mir-155 in HeLa cells, elevating the miRNA from x18 to x90 caused a drop in the average retention of its targets, a trend that is even more prominent at higher overexpression levels. A minor, but consistent increase in retention of non-target genes (Figure 4C, gray color) is consistently observed.

**Figure 5.**
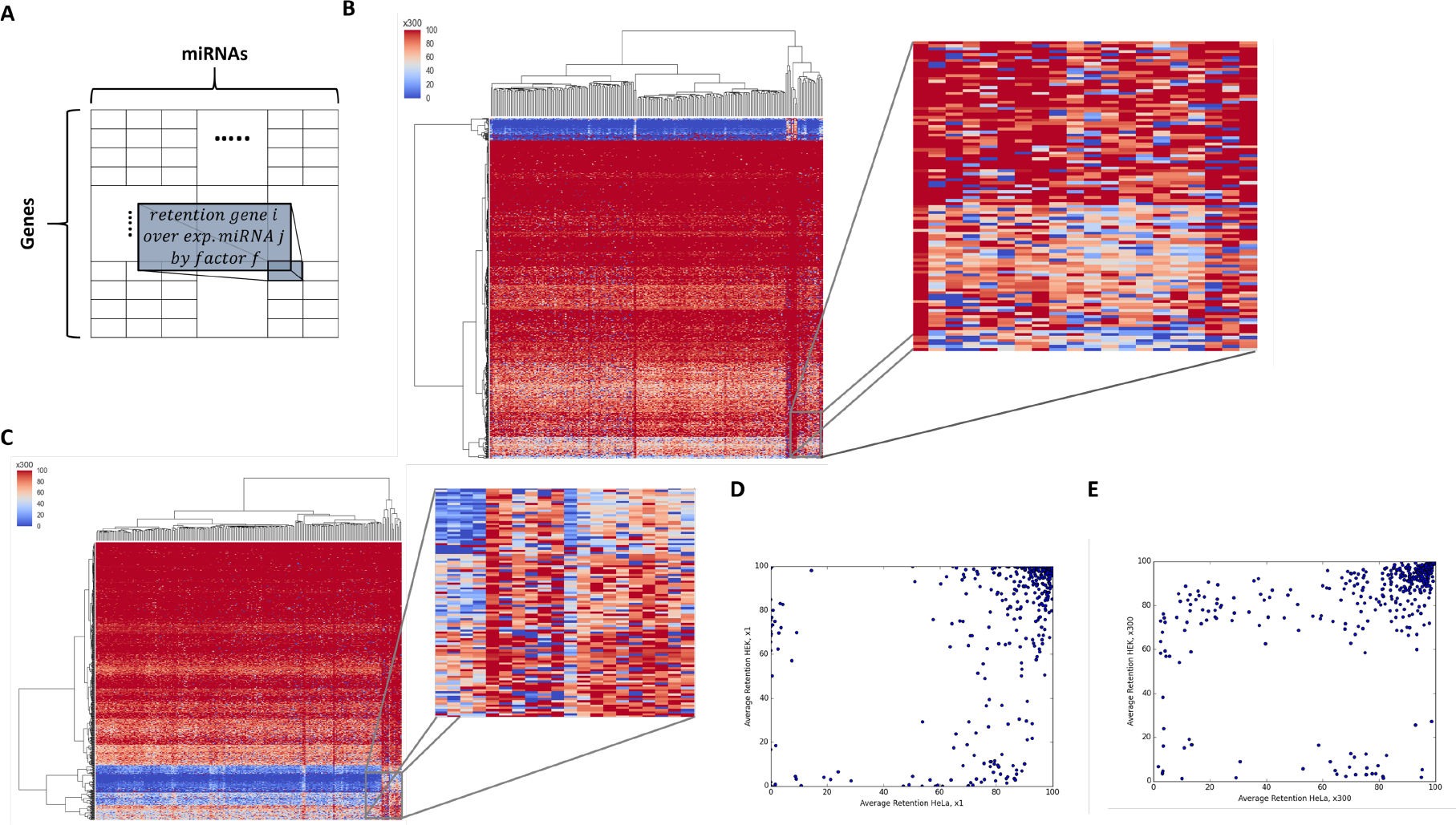
miRNA overexpression matrices. **(A)** The columns stand for the different miRNAs overexpressed by factor f, and the rows stand for the different genes**. (B)** Heatmap of the range of the retention for genes that were overexpressed at a factor x300. Each row is associated with a specific gene. The clustering is performed by the row (i.e. genes). The matrix includes 248 expressed miRNAs in HeLa cells. **(C)** Zoom-in of a small section of the heatmap of the range of the retention for genes that were overexpressed at a factor x300. Each row is associated with the retention by overexpressing anty of the miRNAs. The clustering is performed by the row (i.e. genes). **(D)** HeLa and HEK-293 average final retention comparison in a control setting. Each point stands for each gene. **(E)** HeLa and HEK-293 average final retention comparison. Each point stands for each gene in all 248-overexpression conditions (each row in the heatmap presented in **B, C**) using over expression factor x300.

### A unified pattern of mRNA retention is associated with overexpression of miRNAs

To determine whether the composition and stoichiometry of miRNAs and mRNAs dictate miRNA regulatory behavior, we performed exhaustive and systematic manipulations of all cellular miRNAs (For miRNA profiles for different cell lines, see Supplemental dataset S5). We first clustered individual miRNAs according to their families. For HeLa cells, 248 miRNAs were compiled and match their representation in the miRNA-MBS TargetScan prediction table (see Materials and Methods). We multiply the basal abundance (x1) of each of miRNA families by a factor (*f*) to get a final retention table of 773 genes (rows) whose initial expression exceeds a pre-determined threshold, and 248 miRNA (columns). As each miRNA family was overexpressed by the tested factor (*f*), we obtained a series of matrices for each factor (x3, x9, x18, x90, x300 and x1000). Thus, matrix M*f*_ij_ is the final retention of gene *i* after 100k iterations of COMICS for the overexpressed experiment of miRNA *j* (Figure 5A, Supplemental Dataset S6). For unexpressed miRNA, a minimum level of expression is assigned to the miRNA (x1 level, see Materials and Methods).

Inspecting M*f*_ij_ for each overexpression condition reveals the presence of a substantial set of genes that are characterized by high retention >85% for ≥90% of the tested miRNAs (i.e., the high retention criterion satisfied by at least 225 of all 248 tested miRNA families). We refer to this set as cross-miRNAs stable genes. The number of stable genes is 185 genes for HeLa, 176 genes for HEK-293 and 124 genes for MCF-7. A full detailed analysis of cross-miRNAs stable genes is available in Supplemental Dataset S7. These surprising observations imply that a set of genes in each cell type is resistant to gene expression attenuation regardless of the identity of the overexpressed miRNA. Such coordinated, concerted action of miRNAs seems to be valid for any tested cell type, and to the best of our knowledge was not described previously.

The matrix M*f* also reveals a small defined gene set that is highly sensitive to miRNA regulation. Specifically, these are genes with a retention rate below 50% for ≥90% of the tested miRNAs among all overexpression experiments. These genes are referred to as cross-miRNAs sensitive genes. We report on these sensitive genes for HeLa (23 genes), HEK-293 (34 genes) and MCF-7 (22 genes). For a full detailed analysis of cross-miRNAs sensitive genes see Supplemental Dataset S8. These results imply that a small set of genes in each cell type is prone to regulation by (almost) any overexpressed miRNA. Therefore, attenuation in gene expression is expected for a small set of genes regardless of the actual miRNAs’ composition.

For illustration, the matrices [Mfij (x300)] for HeLa (Figure 5B) and HEK-293 (Figure 5C) are colored to indicate genes with high (red) and low retention (blue) levels. The matrices represent clustering by genes and miRNAs and the clustering dendrogram emphasizes the appearance of a strong signal of sensitive genes (blue rows) in both cell types. Still, a remarkable richness is associated with each of the retention patterns (Figure 5C, zoom in, Mf_ij_ x300). The few miRNAs that are naturally clustered by their similar profile according to their columns will not be further discussed.

We then tested whether the characteristic of the retention profiles is invariable among cell types. Figures 5D-5E compare the average retention observed for each of the shared genes from HeLa and HEK-293. A large difference is observed in the distribution of genes for HeLa and HEK29 when profiles in Mf_ij_ (x1) and Mf_ij_ (x300) are compared. This global view suggests that each manipulated cell converges to its unique cell state, potentially reflects the sensitivity of the analyzed genes to the unique cell composition of miRNAs.

To better characterize the cross-miRNA stable genes, we tested the correspondence of the genes and their cellular function in each of the analyzed cells. Figure 6A shows the unified pattern for Mf_ij_ that was found in HeLa, HEK-293, and MCF-7 for the cross-miRNAs stable and sensitive gene sets. Note that the analysis is limited by the subset of genes common to all three cell types, and express above a predetermined threshold (>0.04% of expressing mRNA molecules). From 78 (MCF-7), 102 (HeLa) and 110 genes (HEK-293) cross-miRNAs stable genes, 48 genes are common to all three cell types. The overlap of this number of genes is very significant (Chi-Square test p-value = 1.35e-08, Figure 6A). It argues that stable genes are immune to miRNA regulation under a wide range of overexpression settings and across cell types. A phenomenon that corroborates the strong, concerted action of miRNAs in each cell types. The list of shared 48 genes is shown in Supplemental Dataset S9.

**Figure 6:**
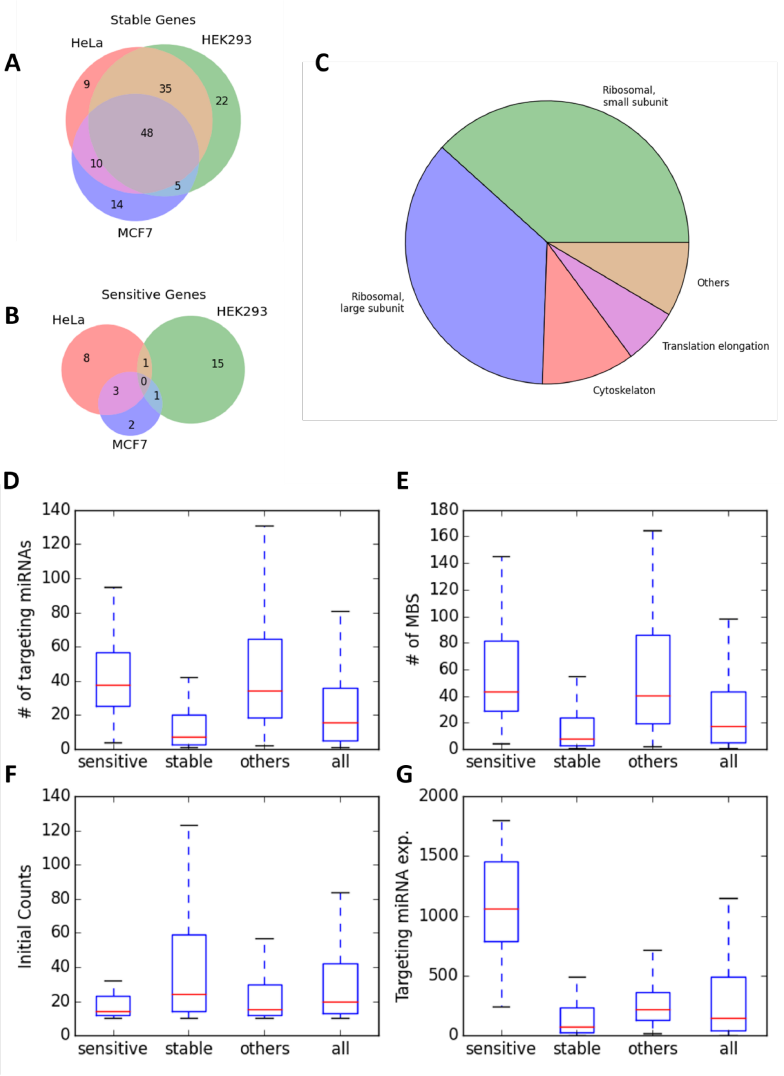
Comparison of sensitive and stable gene sets in different cell types. **(A)** Overlap of the cross-miRNA stable genes in HeLa, HEK-293 and MCF7 cells. Only genes that are expressed in at least two cells are listed. The gene list of the stable genes is available in Supplemental Dataset S7. **(B)** Overlap of the cross-miRNA sensitive genes in HeLa, HEK-293 and MCF7 cells. Only genes that are expressed in at least two cells are listed. The gene list of the stable genes is available in Supplemental Dataset S8. **(C)** The 48 stable genes shared by HeLa, HEK-293 and MCF-7 cells according to their functional annotations: (i) small ribosomal subunit (18 genes), (ii) large ribosomal subunit (17 genes), (iii) cytoskeleton (5 genes), (iv) translation elongation (3 genes) (v) 4 additional genes. For detailed list see Supplemental Dataset S9. **(D-G)** Comparison of the number of targeting miRNA of sensitive genes, stable genes, and other (not sensitive and not stable in HeLa cells). **(D)** Statistics of the comparisons are significant for the comparison of stable genes set and both sensitive and other gene sets (t-test p-values of 7.53e-11 and 6.48e-21, respectively). No significant difference between sensitive and other gene sets. **(E)** Comparison of the number of MBS of sensitive genes in HeLa cells. Statistics of the comparisons are significant for the comparison of stable genes set and both sensitive and other gene sets (t-test p-values of 2.07e-9 and 5.6e-15, respectively). No significant difference between sensitive and other gene sets. **(F)** Comparison of the initial abundance of sensitive genes. Statistics of the comparisons are significant for the comparison of stable genes set and both sensitive and other gene sets (t-test p-values of 0.017 and 0.015, respectively), and no significant difference between sensitive and other gene sets. **(G)** Comparison of the of the average expression of the targeting miRNAs of each gene in HeLa cells. Significant differences between all three gene sets were found (t-test p-values 2.52e-22, 3.07e-9 and 8.7e-11 for the comparison of stable-sensitive, stable-other and sensitive-other, respectively). Full statistics are shown in Supplemental Table S1.

A similar analysis performed for the cross-miRNAs sensitive genes (Supplemental Dataset S8) resulted in an opposite trend. Not only that this gene set is far smaller (Figure 6B), more importantly, no shared genes are detected for the three cell types. Inspecting annotation enrichment reveals that there is no functional coherence for these cell-specific genes.

### Cross-miRNAs stable genes are enriched in translation machinery genes

We applied annotation enrichment tools (see Materials and Methods) to the set of cross-miRNA stable genes from HeLa cells (185 stable genes, Supplemental Dataset S7). We found that these genes are extremely enriched in terms associated with numerous aspects of translation, including translational elongation (GO:0006414), mitochondrial translation (GO:0032543), SRP-dependent co-translational protein targeting to membrane, translational termination (GO:0006415) and more. Combining coherent annotations yield statistically strong enrichment signal (DAVID tool, corrected FDR p-value = 1e-77 to 1e-53) for ribosome structure, elongation machinery, and translational fidelity. Notably, enrichment of annotations associated with protein translation and translation machinery applies for the cross-miRNA stable gene lists from all three cell types (Supplemental Dataset S10).

Figure 6C focuses on annotation partition for the 48 genes that are common to all three cell lines (Supplemental Dataset S9). The dominant role of translational machinery (Annotation clustering enrichment score ~1e-49) unified annotations of translation elongation and cytosolic ribosome (FDR p-value of 1.18e-67 and 9.36e-60, respectively, Supplemental Dataset S10). Translational machinery component with small and large subunits (35 genes), elongation factors (EIF4A1, EEF1D, and EEF1B2) and nucleolin (NCL) account for 79% of the genes in the list. Many of these genes play a role in ribosome production and its dynamic function.

We conclude that the cross-miRNA stable gene set signifies the translational machinery. Specifically, the translational machinery highlights a functional gene set that is immune to the regulatory layer of miRNAs. This observation applies to all tested cells and proposes an overlooked cellular robustness to miRNA perturbations.

Finally, we tested the properties that characterize genes associated with the cross-miRNA stable and sensitive gene sets with respect to other genes that were included in the simulation process (Supplemental Datasets S7-S8). Four properties were tested: (i) the number of targeting miRNAs (Figure 6D), (ii) the number of MBS (Figure 6E), (iii) the initial expression level (Figure 6F), and (iv) the binding potential according to the expression of the most dominant miRNAs (Figure 6G, Supplemental dataset S5). Features (iii-iv) should be considered as cell specific. We observed that genes belonging to the stable set are characterized by fewer MBS and fewer targeting miRNA relative to other genes (t-test = 6.48E-21 and 5.64E-15, respectively). While the statistics for the initial expression levels of these genes are marginal in all cell types, the most significant differentiating feature between the two gene sets is the targeting potency by the most abundant miRNAs in cells (t-test = 2.52E-22, Figure 6G). For example, The most abundant miRNA expressed in HEK-293 is hsa-mir-7 (25% of total miRNA molecules). While it targets only 3.5% of the stable genes, it can bind to 94% of the cross-miRNA sensitive gene set. Therefore, we conclude that stable genes are inherently resistant to regulation by any of the most abundant miRNAs. Table S1 lists the detailed t-test statistics for all three cell lines.

We claim that the exhaustive and unbiased comparison between the stable and sensitive genes reveals an underlying evolutionary signal of MBS. Despite the great difference in overall miRNA composition of different cell lines, several miRNAs are shared across many cell-types (e.g., hsa-mir-21, hsa-let-7 and hsa-mir-92). In all tested cells, the MBS for these genes are excluded from the genes of the translational machinery. Therefore, cellular systems are intrinsically immune to fluctuations in translation by abundant miRNAs.

## Discussion

### miRNAs stability as a major determinant in cell regulation

Cells’ behavior cannot be trivially predicted from the direct measurements of their miRNA and mRNA composition (Arvey et al., 2010; Landgraf et al., 2007). It is assumed that detailed quantitative considerations of miRNA and mRNA govern the dynamics and the steady state of the gene expressed (Bosson et al., 2014; Hausser and Zavolan, 2014). Nevertheless, the underlying rules for post-transcriptional regulation by miRNAs are still missing (Erhard et al., 2014).

We studied cells response to miRNA regulation under a simplified condition of transcriptional arrest by testing the retention profiles of mRNAs as readout. Upon such condition, the dominant effect of miRNAs is via their influence on mRNA stability rather than on translation repression (Bethune et al., 2012). As most miRNAs are transcribed by RNA PolII from their own promoters, it is essential to account for the effect of transcription arrest on miRNA abundance. The results in Figure 1B show that miRNAs are extremely stable at least during the first 24 hrs of ActD exposure. In all studied systems, stabilization of miRNAs is attributed to a protection by AGO-2 (Winter and Diederichs, 2011). Our results (Figure 1) suggest that despite the ActD treatment, AGO-2 is not limiting in the system. Indeed, it was shown that the number of AGO-2 in cells is insensitive to transcriptional arrest (Olejniczak et al., 2013).

The starting point for COMICS simulation is the gene expression pattern as measured experimentally in different cell types (Figure 1, Supplementary Figures S1-S2). COMICS simulation considers a molecular ratio of 2:1 between miRNAs to mRNAs, under the assumption of AGO-2 in excess. For the probabilistic formulation of cells under varying levels of miRNA overexpression, AGO-2 occupancy is governed by recalculating the miRNA distribution without changing the overall stoichiometry (Figure 4A). Thus, avoiding unaccounted limiting factors. We show that changing the number of molecules, while maintaining the stoichiometry impacts the dynamics, with a minimal effect on the endpoint results (Supplemental Figure S4).

### COMICS is robust towards a broad range of parameters

We tested the reliability of COMICS to accurately reflect the miRNA-mRNA competition in living cells. To this end, we altered extensively the simulator’s operational parameters and assessed the sensitivity and robustness of the results (Supplemental File S1). For example, the sampling of miRNAs and mRNAs was tested by applying alternative sampling modes (Supplemental Figure S5). Overall, we demonstrated the robustness of COMICS by varying a large number of parameters (described in Supplementary File S1, Supplemental Figure S4). Moreover, we validated the dependency of COMICS outcome on the quality of the miRNA-MBS table of interactions (Supplemental Figure S5). This table represents a rich computational-experimental body of knowledge (Agarwal et al., 2015).

There are several extensions to COMICS that we intend to explore in the future. This includes the a synergistic cooperativity of miRNA binding at non-overlapping MBS (Balaga et al., 2012; Friedman et al., 2014). Moreover, by applying an option to rationally induce preselected mRNA, quantitative parameters associated with the ceRNA paradigm can be tested. In general, COMICS framework is attractive for testing unresolved questions and emerging principles of miRNA regulation in vivo (Denzler et al., 2014; Hausser and Zavolan, 2014; Yuan et al., 2015).

### miRNA composition is a major determinant in establishing cell identity

We developed COMICS platform to handle cells that undergo a wide range of miRNA manipulations. Altogether, thousands of simulation processes were completed to test the impact of altering the expression of hundreds of miRNA families (Figure 5). We were able to describe general trends from these simulations that apply to three different cell types. Importantly, each of the tested cell types expresses a different profile of miRNAs, supporting the relevance of specific miRNA profiles to cell identity (He et al., 2012) (Supplemental Figure S5). Indeed, transition between cell types is attributed to the expression of a specific miRNAs (e.g. miR-34a (Bu et al., 2013)). Furthermore, specific profiles of miRNAs are associated with the establishment of malignancy states (Bockmeyer et al., 2011; Volinia et al., 2006).

Two extreme miRNA regulation trends were revealed in this study. Firstly, the robustness of the translation apparatus to miRNAs. Secondly, the extreme lability of a small gene set in each cell type. The identification of a small set of genes (about 3% of the reported genes) that is exceptionally sensitive to miRNA regulation, is surprising as it predicts that gene expression down-regulation will result from altering almost any miRNA (Figure 6B). Remarkably, many of the sensitive genes are nuclear, and play a role in transcription regulation. Over-enrichment in MBS for major miRNAs in cells signify many of these genes (Supplemental Table S1). For example, targeting by hsa-mir-21 is prevalent among the cross-miRNA sensitive genes. The hsa-mir-21 occupies 27% and 32% of total miRNAs in untreated MCF-7 and HeLa cells, respectively (Supplemental Dataset S5). The hsa-mir-21 occupies 58 and 73 MBS among the 22 and 34 sensitive genes identified in MCF-7 and HEK-293, respectively.

We plan to extend the use of COMICS to a large collection of human cell lines (e.g., NCI-60) for assessing the generality of our findings. Specifically, applying COMICS simulations to cells that are signified by metastatic potential is attractive for proposing a specific miRNA manipulation that will lead to the desired outcome.

### Evolution of genes toward the most abundant miRNAs

A relatively large gene set (about 20% of the reported genes) that is exceptionally stable to almost any miRNA manipulations, irrespectively to the actual levels of the miRNA is unexpected (Supplemental Datasets S6, S8). Based on the statistically significant functional overlap among three cell types it is postulated that the 3’-UTR of these stable genes is under a strong evolutionary selection. Indeed, the most significant difference between the features associated with the sensitive and stable sets (p-value = 1.57E-23) concerns the cellular level of the miRNA expression (Figure 6G). In cancer cells, the most abundant miRNA genes (e.g., hsa-mir-21, hsa-mir-30, hsa-mir-15/16) are involved in cancer development. In these instances, a minor change in their expression leads to a transition in cell states (Volinia et al., 2010). In view of our finding, we propose that during evolution time many of the ribosomal proteins and the components of the translation machinery became robust to miRNA regulation. We propose that the translation machinery has evolved to achieve a maximal robustness vis-a-vis miRNA regulation. Genes of the translation components are practically resistant to changes even in the presence of abundant miRNAs.

In summary, the immunity of the translation system to miRNA regulation suggests that it may be part of a global cell strategy (Lopez-Maury et al., 2008). Evidently, there is a fundamental difference between transcription and translation processes. While the transcription system can quickly respond to the needs dictated by changes in the environment, the translational machinery is stable and much less prone to variations. Therefore, the sustaining immunity towards the majority of miRNAs, including the high expressing miRNAs unveils an overlooked design principle in miRNA regulation. The driving force of evolution, acting on targets and their MBS underlying the properties of the miRNA network in many organisms (Berezikov, 2011). It is for the future to investigate whether this evolutionary refinement of 3’-UTR of the translation apparatus is a general trend that dominate the miRNA regulatory network in other organisms.

## Materials and Methods

### Cell culture

Human cell line of HeLa (cervix epithelial, # CCL-2) and HEK-293 (embryonic kidney, # CRL-1573) were purchased from the cell-line collection of ATCC. Cells were cultured at 37^0^C, 5% CO_2_ in Dulbecco’s Modified Eagle Media (DMEM, Sigma), supplemented with 10% FBS (Life Technologies), and 1% antibiotics mixture (Sigma-Aldrich, Cat # P4333). Cells were maintained for 2 weeks and passing and splitting cells was carried out at 70-80% confluence.

### Transcription arrest and miRNA overexpression

Overexpression of miRNAs was performed by transfected HeLa cells and HEK-293 with miRNA expression vectors that are based on the miR-Vec system, under the control of CMV promotor (Origene). Cell transfection was done using Lipofectamine 3000 (Invitrogen) as described by the manufacturer. Cells at 70% to 80% confluency were transfected with 1.5μg purified plasmid DNA containing hsa-mir-155 and hsa-mir-124a (kindly contributed by Noam Shomron, Tel Aviv University). Control empty vector expressing GFP (0.15μg) was mixed with the CMV-miR expressing vectors. Cells were monitored by fluorescent microscopy at 36 and 48 hrs post transfection. The efficiency of cell transfection was >75% of the HeLa cells and ~100% of the HEK-293 according to the GFP expression at 48 hrs post transfection. Transcription inhibition was achieved by adding to cultured HeLa and HEK-293 cells media containing Actinomycin D (ActD, 10 μg/ml in DMSO), or the appropriate control (i.e. DMSO). Cells were treated with ActD (10 μg/mL, Sigma) 24 hrs post-transfection. Cells were cultured in 6-well plates and following treatment were lysed in 1 ml TRIzol (Invitrogen) at the indicated time points (0 hrs, 2 hrs, 8 hrs, 24 hrs). Expression normalization for mRNA and miRNA and quantitation are described in Supplemental File S1. Protocols for RNA and miRNA library preparation, RNA deep sequencing analysis are available in Supplemental File S1.

### Probabilistic based miRNA-mRNA simulator

The simulator input is the number of molecules for the expression profiles of miRNAs (total 50k molecules) and mRNAs (total 25k molecules) in the specific cell type, and a table of miRNA-mRNA interaction prediction extracted from TargetScan (Supplementary File S1). In addition, the simulator, called COMICS (COmpetition of MiRNA Interactions in Cellular Systems) supports a wide set of configurable parameters: (i) the number of total miRNA; (ii) the number of mRNA molecules in the cell; (iii) the number of iterations for completing the run; (iv) the number of iteration interval between miRNA-mRNA binding event and the mRNA removal; (v) a random removal of unbounded mRNAs according to predetermined decay rate of the mRNA as extracted and extrapolated from experimental data of mRNA half-life; (vi) addition of newly transcribed mRNAs during a configurable number of iterations interval; (vii) miRNAs or genes overexpression according to a selected multiplication factor for the degree of overexpression. (vii) incorporation of alternative miRNA-target mapping. It is also possible to activate the simulator by a set of random genes as an initial state of pre-existing iterations prior to the simulation run. The protocol applied for each iteration is described in Supplemental File S1.

Overexpression scheme is based on multiplication of the available miRNA amount by all factors (from x1 to x1000). This addition of miRNA molecules calls for calculating a new miRNA distribution while remaining with the same amount of miRNA in the cell. In case the miRNA had not detected in the native cell, an arbitrary starting minimal amount of 0.01% (the equivalent of 5 molecule/ cell).

### Statistics and Bioinformatics

P-values were calculated using a paired and unpaired t-test, Fisher exact test, Kolmogorov Smirnov (KS) test or Chi-square tests. For testing the correspondence of two sets of different sizes, we have used the Jaccard score (J-score) that is the size of the intersection divided by the size of the union of the sample sets and it is range from 0 to 1 (no correspondence to a complete overlap, respectively). Statistical values are that are based on correlations were performed using standard Python statistical package. For annotation enrichment statistics (Kuleshov et al., 2016) was used. For testing the effect of different background gene lists for the enrichment statistics we applied DAVID (Huang et al., 2007) clustering enrichment score is based on one tail Fisher exact corrected for the number of gene ontology annotations that are used. Enrichment was performed in view of genes that are potential candidates for our analysis and against the set of genes that express with a minimum of 0.02% of the mRNA overall expression.

## Acknowledgments

The study was supported in part by ERC grant 339096 on High Dimensional Combinatorics. We thank Tsiona Eliyau for supporting the experimental data for HeLa and HEK-293. We thank Noam Shomron (Tel Aviv University) for sharing the miRNA expression plasmids used in this study. We thank the members of the Linial’s lab for useful comments and discussions.

## Author contributions

SMA, NL and ML contributed to the experimental design and leading the research for this study. All authors provided a critical review. SMA developed the cell simulator, executed the results and conducted the data modeling. SMA wrote the computational code for the implementation of COMICS platform, SMA, NL and ML, wrote the paper.

## Additional files

### Supplemental Datasets S1-S10

- The mapped mRNAs and miRNAs for HeLa and HEK-293 cells are listed in Supplemental Dataset S1 and S2, respectively.
- The output of the % retention along 1M COMICS iterations at a 10k resolution from HeLa as input is shown in Supplemental Dataset S3.
- The output of the % retention along 100k COMICS iterations at a 1k resolution is shown for HeLa and HEK-293 cells in Supplemental Dataset S4.
- The lists of miRNA expression levels for 3 cells as input for COMICS are in Supplemental Dataset S5.
- The Matrix of the miRNA retention levels for 248 miRNAs and 773 genes are in Supplemental Dataset S6.
- Lists of the stable genes for the three cells are in Supplemental Dataset S7.
- Lists of the sensitive genes for the three cells are in Supplemental Dataset S8.
- The results from enrichment tests for functional annotation for stable genes that are shared among 3 cell lines (48 genes) are in Supplemental Dataset S9.
- The results from enrichment tests for functional annotation for stable genes for 3 cell lines are in Supplemental Dataset S10.

### Supplemental File S1 (text)

- Supplemental File S1 presents A. Extended Materials and methods (A1-A6); B. The results for the alteration of the operational parameters of COMICS. C. Figure legends Figure S1-Figure S6.

### Supplemental Table

- Table S1 summarizes the statistics of characteristics for sensitive and stable sets for 3 cell lines.

### Supplemental Figures S1-S6

- Supplemental Figure S1 and Figure S2 show the time point correlations of the expression of miRNAs and mRNAs from HeLa and HEK-293, respectively.
- Supplemental Figure S3 shows the retention profile for hsa-mir-124a in HeLa cells.
- Supplemental Figure S4 shows the results of alteration of COMICS parameters.
- Supplemental Figure S5 shows the results of repeated runs and randomization protocol for the probabilistic miRNA-mRNA interaction table.
- Supplemental Figure S6 shows the heatmap of the most abundant miRNA in 3 cell lines and the result of shuffling between them.

